# Immunoinformatic design of multi epitopes peptide-based universal cancer vaccine using matrix metalloproteinase-9 protein as a target

**DOI:** 10.1101/2020.02.16.951319

**Authors:** Abdelrahman H. Abdelmoneim, Mujahed I. Mustafa, Miyssa I. Abdelmageed, Naseem S. Murshed, Enas A. Dawoud, Enas M. Ahmed, Rahma M. Kamal eldin, Nafisa M. Elfadol, Anfal Osama M. Sati, Abdelrafie M. Makhawi

**Affiliations:** Faculty of Medicine, Alneelain University, Khartoum, Sudan; Department of Biotechnology, University of Bahri, Khartoum, Sudan; Faculty of Pharmacy, University of Khartoum, Khartoum, Sudan; Department of Microbiology, International University of Africa, Khartoum, Sudan; Faculty of Medical laboratory sciences, Razi University, Khartoum, Sudan; Faculty of Medicine, Karary University, Khartoum, Sudan; National University Biomedical Research Institute, National University, Khartoum, Sudan

**Keywords:** Immunoinformatics, Peptides, Universal Cancer Vaccine, IEDB, matrix metalloproteinase-9

## Abstract

**Background:** Cancer remains a major public health hazard despite the extensive research over the years on cancer diagnostic and treatment, this is mainly due to the complex pathophysiology and genetic makeup of cancer. A new approach toward cancer treatment is the use of cancer vaccine, yet the different molecular bases of cancers reduce the effectiveness of this approach. In this work we aim to use matrix metalloproteinase-9 protein (MMP9) which is essential molecule in the survival and metastasis of all type of cancer as a target for universal cancer vaccine design.

**Method:** reference sequence of matrix metalloproteinase-9 protein was obtained from NCBI databases along with the related sequence, which is then checked for conservation using BioEdit, furthermore the B cell and T cell related peptide were analyzed using IEDB website. The best candidate peptide were then visualized using chimera software.

**Result:** Three Peptides found to be good candidate for interactions with B cells (SLPE, RLYT, and PALPR), while ten peptides found as a good target for interactions with MHC1 (YRYGYTRVA, YGYTRVAEM, YLYRYGYTR, WRFDVKAQM, ALWSAVTPL, LLLQKQLSL, LIADKWPAL, KLFGFCPTR, MYPMYRFTE, FLIADKWPA) with world combined coverage of 94.77%. In addition, ten peptides were also found as a good candidates for interactions with MHC2 (KMLLFSGRRLWRFDV, GRGKMLLFSGRRLWR, RGKMLLFSGRRLWRF, GKMLLFSGRRLWRFD, TFTRVYSRDADIVIQ, AVIDDAFARAFALWS, FARAFALWSAVTPLT, MLLFSGRRLWRFDVK, GNQLYLFKDGKYWRF, NQLYLFKDGKYWRFS), with world combined coverage of 90.67%.

**CONCLUSION:** 23 peptide-based vaccine was designed for use as a universal cancer vaccine which has a high world population coverage for MHC1(94.77%) and MHC2 (90.67%) related alleles.

## 1. Introduction

Cancer is a chronic disease with varying degree of manifestations characterized by abnormal cell division and high mortality and morbidity rates, in addition it is considered the second leading cause of death worldwide with an estimated 8 million death annually and 18 million new case per year.(1-5) although cancer survival rate increased dramatically over the years in many countries, yet more work are needed to improve the prognosis of patients suffering from this depilating disease.(6, 7) Cancer can be either common or rare depending on the number of cases. Usually any cancer with a number of cases below 6 in 100000 annually is considered rare type.(8) All types of cancers cause common clinical symptoms including weight-loss, fatigue, pain, nausea, vomiting and constipation, etc.(9) Additional symptoms are specific to the site of malignancy.(10) Breast cancer is the most common site of cancer in women worldwide.(11) Lung cancer is the second most common cancer in both men and women while colorectal is the third most common type cancer in both sexes. (12)

The classical management for cancer composed of chemotherapy, radiotherapy and surgery. Recently Immunotherapy started to shows very promising result in field of cancer management. Different type of immunotherapy are available including cytokine therapy, checkpoint inhibitors, monoclonal antibodies and therapeutic vaccines.(13, 14) With the latter being the center of many researches in the recent years, cumulating in the approving of Sipuleucel-T By the U.S. Food and Drug Administration (FDA) as the first dendritic cell based therapeutic vaccine for prostate cancer in 2010.(15, 16) vaccine can either be used alone or in combination with other modalities like immune checkpoint inhibitors (ICIs) which shows better results when compared with vaccine monotherapy.(17)

Most drug company invests in the common types of cancer like breast and colorectal cancer for financial and cost effective reasons leaving the rare type of cancer like chondrosarcoma with no effective mode of treatment.(18-20) This call for a new strategy for management of cancer as a whole rather than treating different types individually. Furthermore, although many trials of therapeutic and preventive drugs to cancer has been carried out throughout the years, yet none of them succeeded to obtain a positive response from all types of cancers, moreover, the FDA approved none as universal cancer vaccine agent.(21, 22) The reason for this failure is the fact that no effective intervention is available that covers all the pathological pathways of all types of cancer caused by the massive tumor heterogeneity.(23)

There are approximately 28 subtypes of matrix metalloproteinase (MMPs). They are responsible of maintaining the components of the extracellular matrix (ECM), and controlling metastasis of cancerous cells. *Matrix metalloproteinases-9 (MMP-9)* gene (MIM 120361) is located at chromosome 20q12-13. It encodes Matrix metalloproteinase-9 protein, a major enzyme for the degradation and absorption of ECM, signaling of wound healing, bone resorption, angiogenesis and plays a crucial role in various types of cancer progression. Such as breast cancer, colorectal cancer, papillary thyroid cancer, urinary cancers, lymphomas and gastric cancer.(24-27) In addition, MMP-9 is considered as a biomarker as its level increases in inflammatory diseases, i.e., Rheumatoid arthritis, asthmas, and abdominal aortic aneurysm.(28, 29) This study aims to predict a safe multi-epitope based anti-cancer vaccine using human MMP-9 protein as a target.

MMP-9 protein was tested in mouse model before for several therapeutic purpose,(30, 31) but to the best of our knowledge, this is the first immunoinformatics study to use human MMP9 protein as an immunogenic target for a peptide-based vaccine design for all cancer types. Peptide based vaccines are safe, stable with minimal exposure to hazardous or allergenic substances, cheaper and less laboring.(32, 33) In Addition, reverse vaccinology techniques have proven its efficiency in general and in cancer field in particular, as many of the designed vaccines has made it into clinical trials.(34-36)

## 2. Materials and Methods

### 2.1 Sequences retrieval

The whole protein sequence of the MM9 in FASTA format, accessed on November 3,2019 (NP-004985.2) was retrieved from NCBI database (https://www.ncbi.nlm.nih.gov/protein).(37) On the other hand, the candidate epitopes were analyzed by IEDB analysis tools (http://www.iedb.org/).(38)

### 2.2 Identification of conserved regions

Sequences were aligned to identify the conserved regions using BioEdit tool package version 7.2.5. Furthermore, epitope conservancy investigation for each epitopes was predicted by IEDB analysis resource. (http://tools.immuneepitope.org/tools/conservancy).(39)

### 2.3 B-cell epitope prediction

B cell epitope is the part of a vaccine which interacts with B cell receptor. The epitopes of interest were investigated using numerous B-cell prediction algorisms to identify the antigenicity, flexibility, hydrophilicity and surface accessibility; the selection of highest immunogenic protein sequence is a prerequisite for epitope-based peptide vaccine design.(40)

#### 2.3.1 Prediction of linear B-cell epitopes

BepiPred from immune epitope database (http://toolsiedb.ofg/bcell/) [28] was used as linear B-cell epitopes prediction from the conserved region with a default threshold value of 0.35.all epitopes with values less than this threshold were chosen.

#### 2.3.2 Prediction of surface accessibility

By using Emini surface accessibility prediction tool of the immune epitope database (IEDB) (http://tools.immuneepitope.org/tools/bcell/iedb). [29] The surface accessible epitopes were predicted from the conserved region holding the default threshold value 1.000. we selected all peptides with values less than this threshold for further analysis.

#### 2.3.3 Prediction of epitopes antigenicity

The kolaskar and tongaonker antigenicity method was used to determine the antigenic sites with a default threshold value of 1.028 It is available at (http://tools.immuneepitope.org/bcell/),[30] all antigenic peptide with values less than the threshold were selected for further analysis.

### 2.4 T cell epitope prediction

#### 2.4.1 MHC class I binding predictions

Peptide binding to MHC class I molecules was analyzed by the IEDB MHC I prediction tool (http://tools.iedb.org/mhc1.); based on Artificial Neural Network (ANN) .this tool has different algorithms to show the ability of the submitted sequence to bind to a specific MHC class 1 molecule.(38, 41-43) All epitope lengths were set as 9 amino acids, all conserved epitopes that bind to alleles at score equal or less than 100 half-maximal inhibitory concentration (IC50) is selected for further analysis.(44)

#### 2.4.2 MHC class II binding predictions

Peptide binding to MHC class II molecules was analyzed by the IEDB MHC II prediction tool (http://tools.iedb.org/mhcII) (45, 46) the three major types of HLA alleles (DR, DQ and DP) were chosen for analysis based on NN-align, and all peptide with IC50 less than 500 were chosen for further workout.

### 2.5 Population coverage

Diverse HLA alleles are very distributable world-wide population. Hence, the HLA-alleles distribution among the world population is critical for effective multi-epitope vaccine design. Predicted population coverage analysis of above-mentioned MHC class I and II epitopes were done by using IEDB tool against world-wide population. (http://tools.iedb.org/tools/population/iedb_input).(47)

### 2.6 3D Modeling

The full 3D structure of MM9 was generated by RaptorX (http://raptorx.uchicago.edu/).(48) The accession number of the reference sequence is NP-004985.2 then UCSF Chimera was used to visualize the position of suggested peptides.(49) the modeling was then verified using Ramachandran plot from the model panel in chimera software.

### 2.7 Allergenicity test

AllerTOP v. 2.0 was used to check the allergenicity of MMP9 protein. The method is based on auto cross covariance (ACC) transformation of protein sequences into uniform equal-length vectors. It depends on five E descriptors of amino acids (amino acid hydrophobicity, molecular size, helix-forming propensity, relative abundance of amino acids, and β-strand forming propensity). The proteins are classified depending on training sets containing different known and unknown allergens.(50) It is available at https://www.ddg-pharmfac.net/AllerTOP/index.html.

### 2.8 Antigenicity test

VaxiJen v. 2.0 was used to check the antigenicity of the chosen peptides with a cut point of 0.5. It is an alignment-free approach to in silico antigen identification, based on the auto cross covariance (ACC) method. It has a prediction accuracy ranging from 69% to 97 %.(51) The software is available at http://www.ddg-pharmfac.net/vaxijen/VaxiJen/VaxiJen.html.

### 2.9 Prediction of interferon-gamma inducing epitopes

IFNepitope was used to predict the MHC2 related peptides with a capacity to activate interferon gamma cytokine productions. It is based on two models: Motif based and SVM based models.(52)

The software is available at http://crdd.osdd.net/raghava/ifnepitope/index.php.

## 3. Results

### Prediction of B cell epitope

The metalloproteinase-9 protein (MMP-9) reference sequence was examined by using Bepipred linear epitope prediction, Kolaskar and Tongaonkar antigenicity and Emini surface accessibility methods in IEDB, to confine the probability of specific regions in the protein to bind to B cell receptor, being in the surface and immunogenic, respectively.

In bepipred linear epitope prediction method which was the first method used to determine the B cell epitopes, the average binder’s score of MMP-9 protein to B cell was 0.321, with a minimum score of - 0.006 and a maximum score of 3.227, Basically all values that equal or greater than the default threshold 0.350 were potentially linear epitopes. as shown in (table 1 and figure 1).

**Table1:**
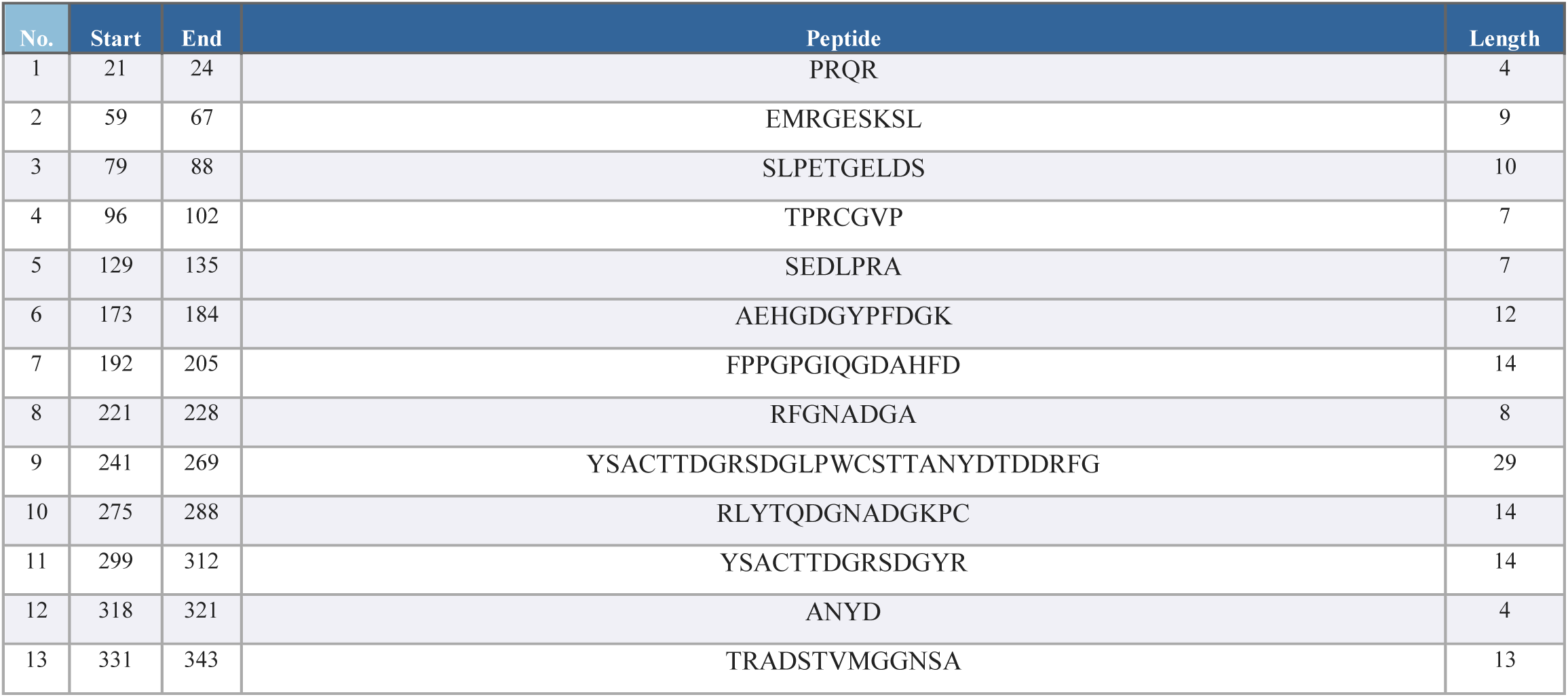
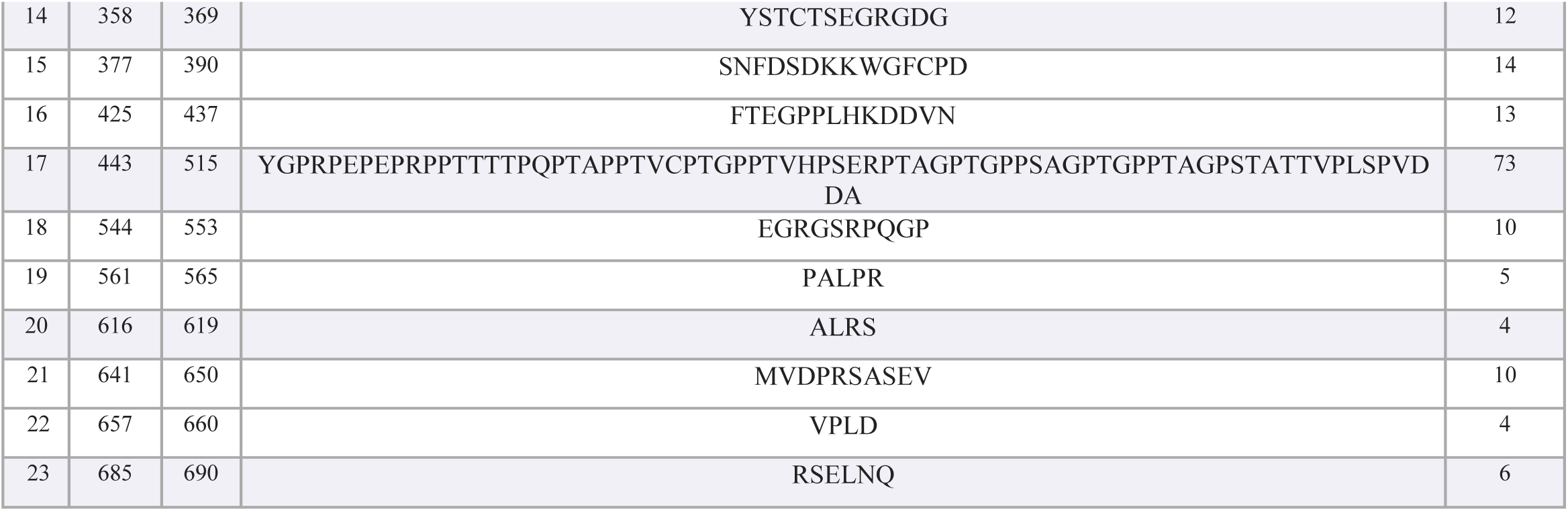
describe the predicted antigenic B cell epitopes, 23 antigenic sites were identified from MMP-9 protein

**Figure 1:**
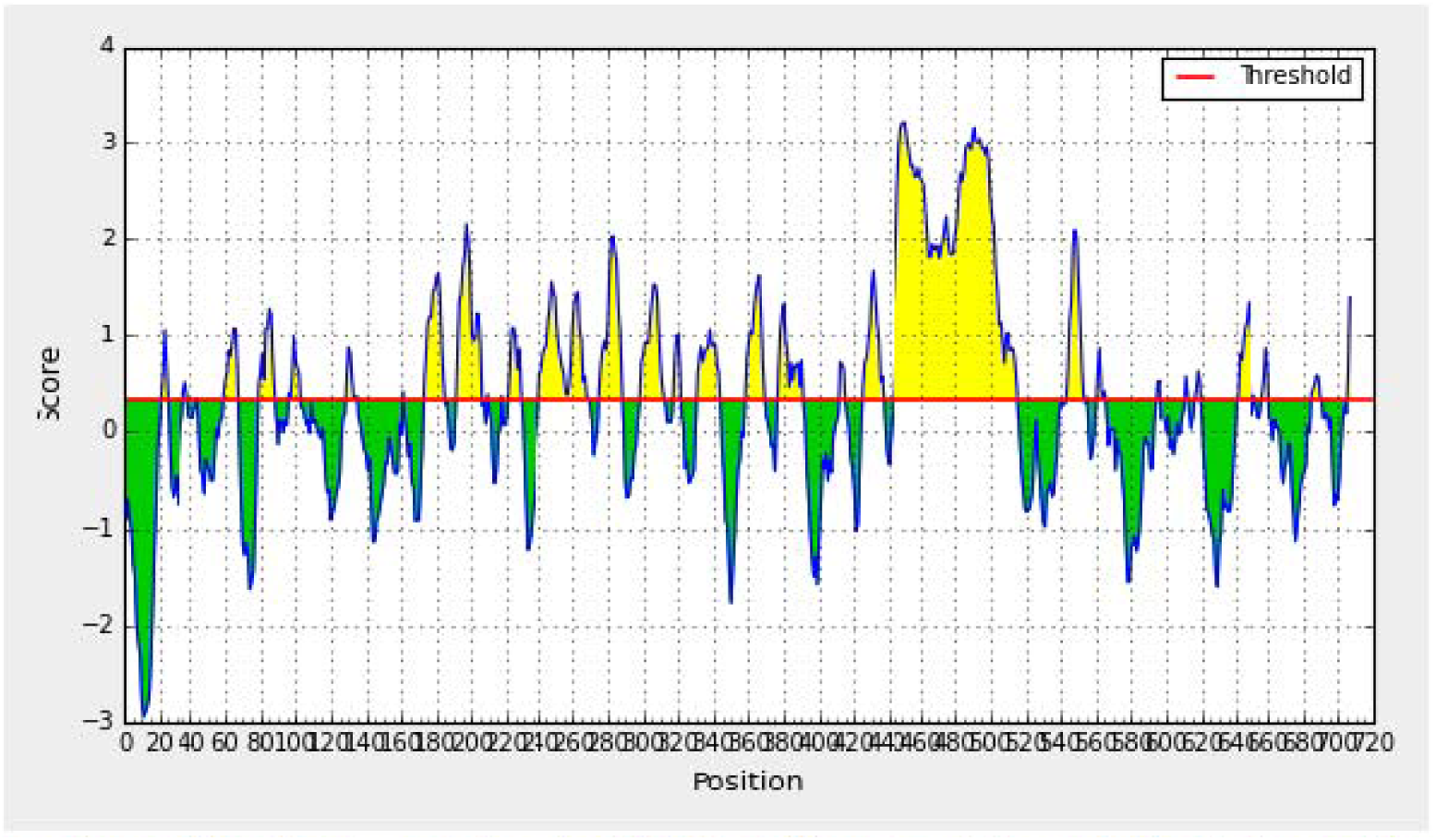
Bepipred linear epitope prediction, the yellow peaks above the red line (Threshold) are proposed to be a part of B cell epitopes and the green peaks are not a part.

In Emini surface accessibility prediction, for an efficient B-cell epitope the average surface accessibility areas of the protein was scored as 1.000, with a maximum of 6.034 and a minimum of 0.031, so all the values equal or greater than the default threshold 1.000 were potentially in the surface. As well as, Kolaskar and Tongaonkar antigenicity prediction’s the average of antigenicity was 1.028, with a maximum of 1.307 and minimum of 0.869; all values equal to or greater than the default threshold 1.028 were potential antigenic determinants. The results of conserved predicted B cell epitopes that passed both Emini & kolaskar tests are shown in (table 2 and figure 2&3).

**Table 2:** List of conserved B cell epitopes with their surface accessibility and antigenicity scores.

| Start | End | Peptide | Length | Conservation | EMINI score | Prediction | KOLSKAR score | Prediction |
| --- | --- | --- | --- | --- | --- | --- | --- | --- |
| 79 | 82 | SLPE | 4 | C | 1.079 | P | 1.044 | P |
| 275 | 278 | RLYT | 4 | C | 1.331 | P | 1.048 | P |
| 561 | 565 | PALPR | 5 | C | 1.104 | P | 1.063 | P |

**Figure 2:**
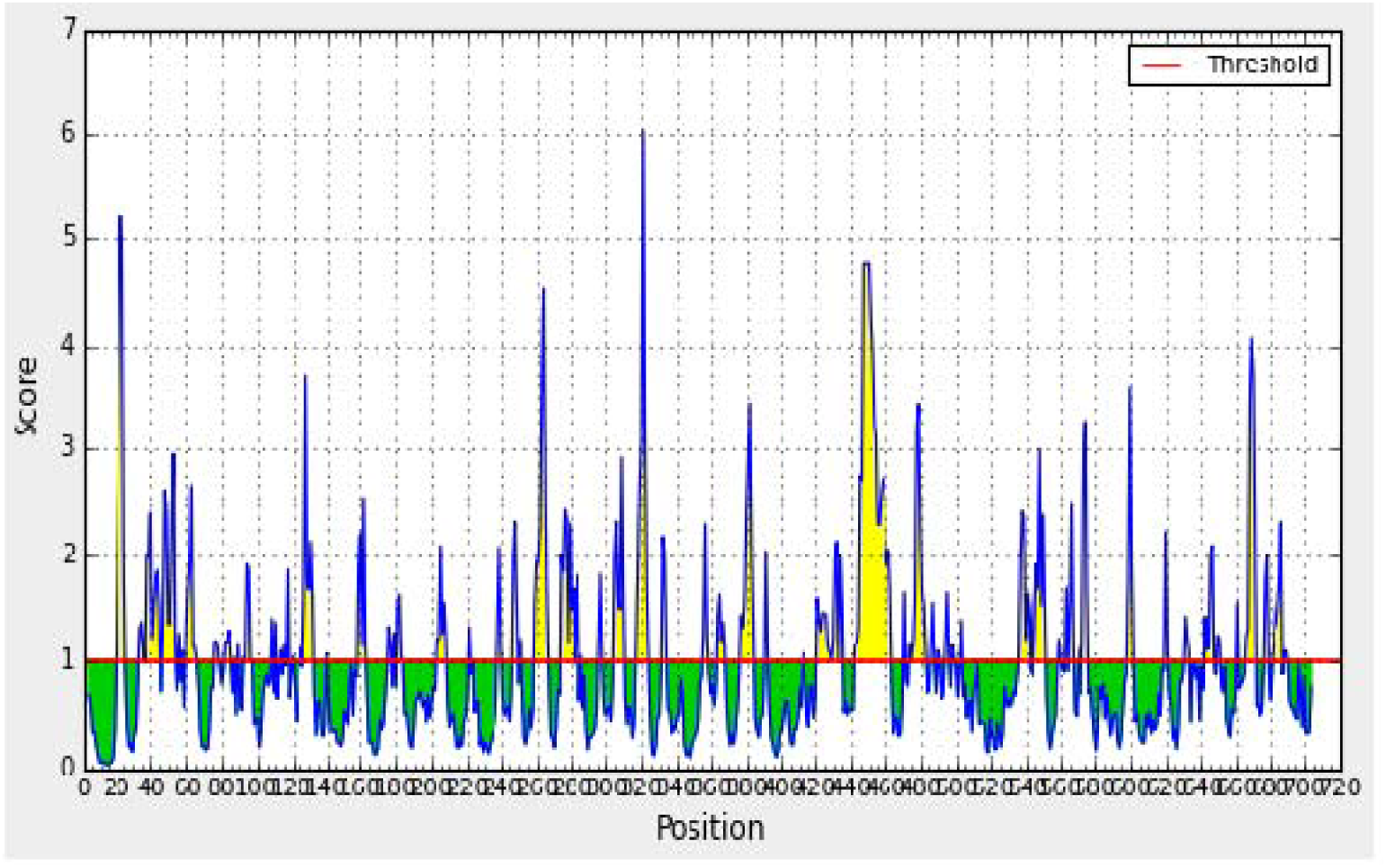
Emini Surface Accessibility Prediction, Yellow peaks above the red line(Threshold) are proposed to be a part of B cell epitopes while green peaks are not a part.

**Figure 3:**
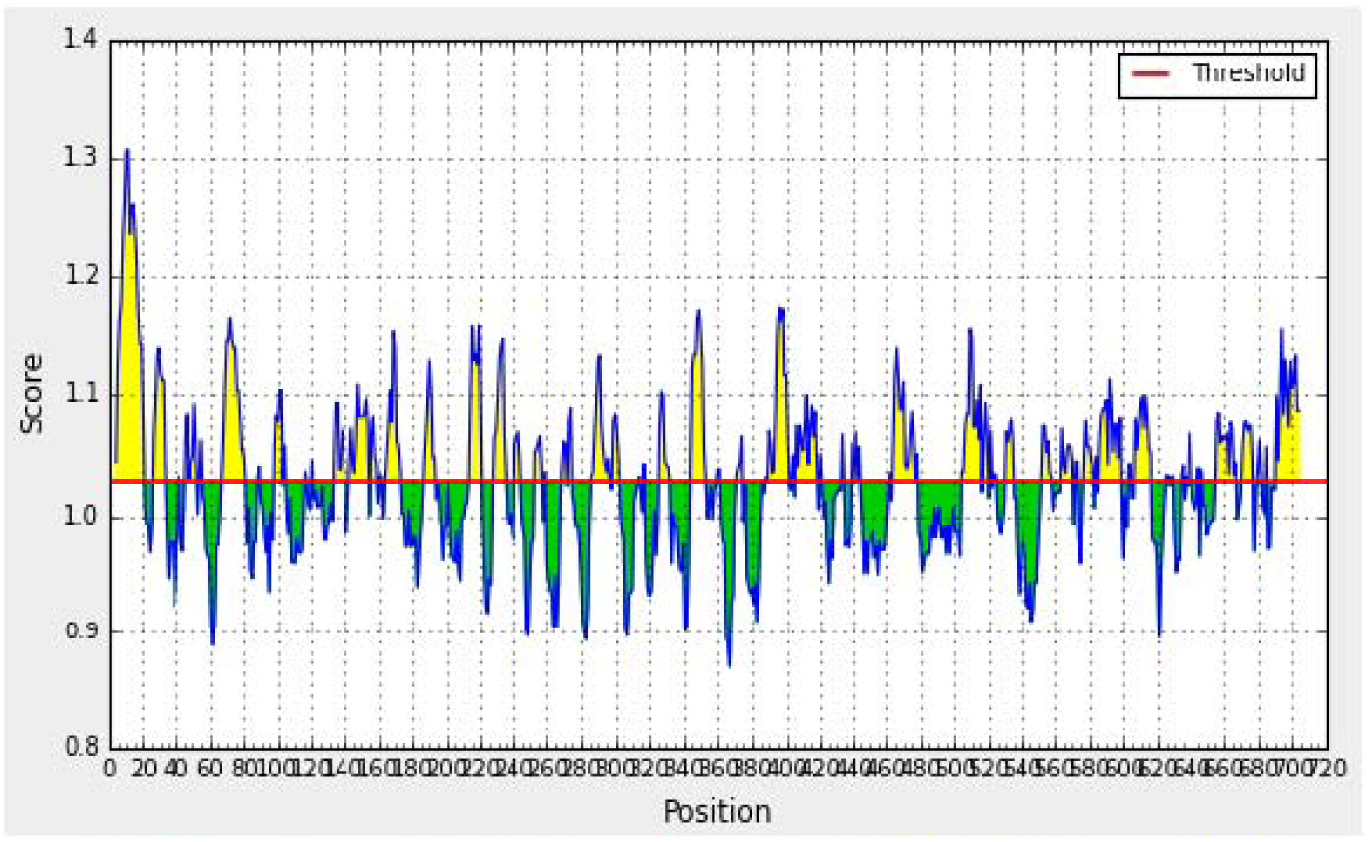
Kolaskar and Tongaonkar Antigenicity Prediction, Yellow peaks above the red line(Threshold) are proposed to be a pan of B cell epitope while green peaks are not a part.

### Prediction of T cell epitope

#### MHC-I binding profile prediction for T cytotoxic cell conserved epitopes

227 epitopes were anticipated to interact with different MHC-1 alleles. The core epitopes (YRYGYTRVA/ YGYTRVAEM) were noticed to be the dominant binders with 7 alleles for each(HLA-C*06:02,HLA-C*07:01,HLA-C*12:03,HLA-C*14:02,HLA-B*27:05,HLA-C*07:02,HLA-B*39:01/HLA-C*03:03,HLA-C*14:02,HLA-C*12:03,HLA-C*06:02,HLA-B*08:01,HLA-C*15:02,HLA-C*07:01). Followed by (YLYRYGYTR,WRFDVKAQM, ALWSAVTPL, LLLQKQLSL, LIADKWPAL, KLFGFCPTR, MYPMYRFTE, FLIADKWPA) which binds with (8, 5, 5, 3, 3, 5, 1, 3) alleles. (Table 3).

**Table 3:** List of most promising epitopes that had a good binding affinity with MHC-I alleles in terms of IC50.

| Peptide | start | end | No. of MHC-I allele hits |
| --- | --- | --- | --- |
| YRYGYTRVA | 50 | 58 | 7 |
| YGYTRVAEM | 52 | 60 | 7 |
| YLYRYGYTR | 48 | 56 | 8 |
| WRFDVKAQM | 633 | 641 | 5 |
| ALWSAVTPL | 146 | 154 | 5 |
| LLLQKQLSL | 72 | 80 | 3 |
| LIADKWPAL | 555 | 563 | 3 |
| KLFGFCPTR | 324 | 332 | 5 |
| MYPMYRFTE | 419 | 427 | 1 |
| FLIADKWPA | 554 | 562 | 3 |

#### MHC-II binding profile prediction for T helper cell conserved epitopes

479 conserved predicted epitopes were found to interact with MHC-II alleles. The core epitopes (KMLLFSGRRLWRFDV) is thought to be the top binder as it interacts with 18 alleles; (HLA-DRB1*01:01,HLA-DRB1*03:01,HLA-DRB1*04:02,HLA-DRB1*07:01,HLA-DRB1*04:04,HLA-DRB1*08:01,HLA-DRB1*09:01,HLA-DRB1*13:01,HLA-DRB1*11:01,HLA-DRB1*13:02,HLA-DRB1*12:01,HLA-DRB1*15:01,HLA-DRB1*16:02,HLA-DRB3*01:01,HLA-DRB4*01:03,HLA-DRB5*01:01,HLA-DRB3*03:01,HLA-DRB4*01:01). Followed by (GRGKMLLFSGRRLWR, RGKMLLFSGRRLWRF,GKMLLFSGRRLWRFD,TFTRVYSRDADIVIQ, AVIDDAFARAFALWS,FARAFALWSAVTPLT,MLLFSGRRLWRFDVK, GNQLYLFKDGKYWRF, NQLYLFKDGKYWRFS) which binds to (17, 17, 18, 14, 14, 17, 15, 14, 14) alleles. (Table 4).

**Table 4:**
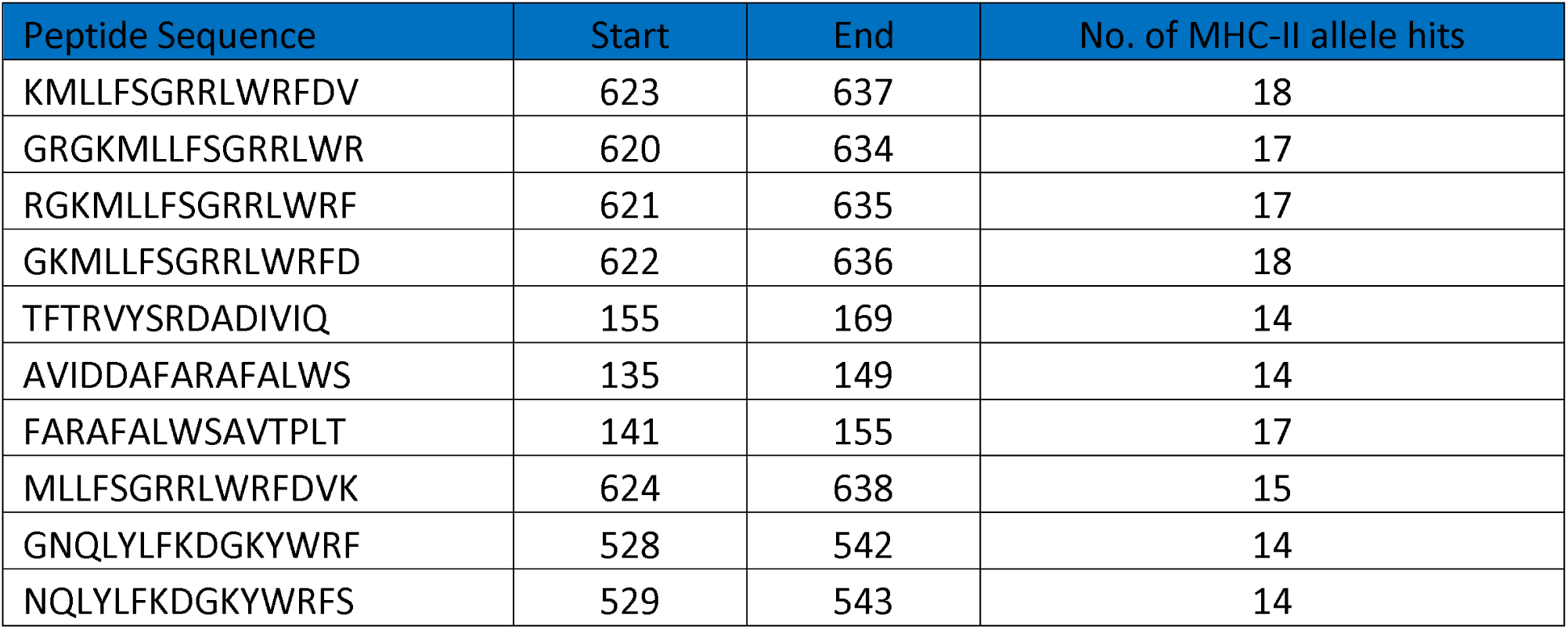
List of the most promising core sequence epitopes that had a strong binding affinity with MHC-II in terms of IC50.

### Population coverage analysis

Population coverage test was performed to detect the world Coverage of all epitopes binds to MHC1 alleles and MHC11 alleles, Exhibit an exceptional coverage with percentages of 94.77%, 90.67%. Respectively, as well as selected most promising epitopes for each test.

#### Population coverage for MHC1

ten epitopes are given to interact with the most frequent MHC class I alleles (YRYGYTRVA, YGYTRVAEM, YLYRYGYTR, WRFDVKAQM, ALWSAVTPL, LLLQKQLSL, LIADKWPAL, KLFGFCPTR, MYPMYRFTE and FLIADKWPA), representing a considerable coverage against the whole world population. The maximum population coverage percentage over these epitopes worldwide was found to be 62.74% for YRYGYTRVA (table 5, figure 4).

**Table 5:**
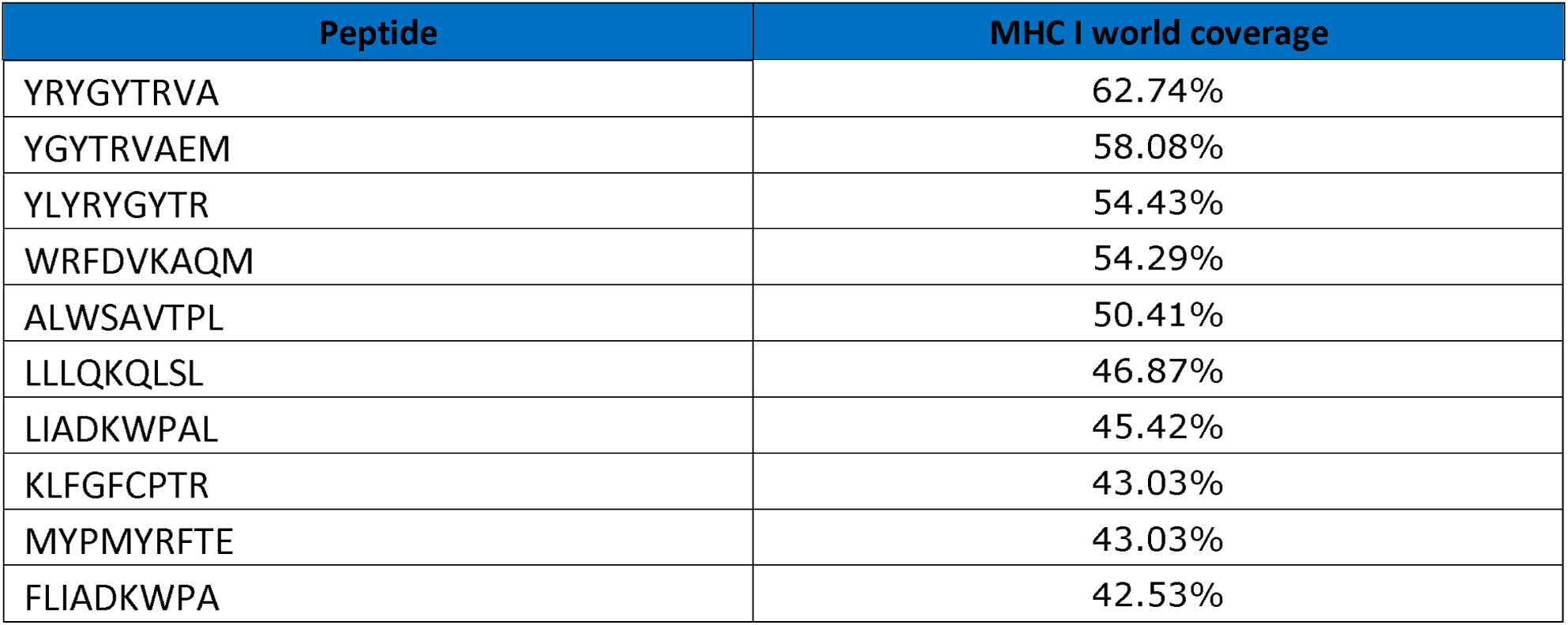
Global Population coverage of promising epitopes binding to MHC class I alleles.

**Figure 4.**
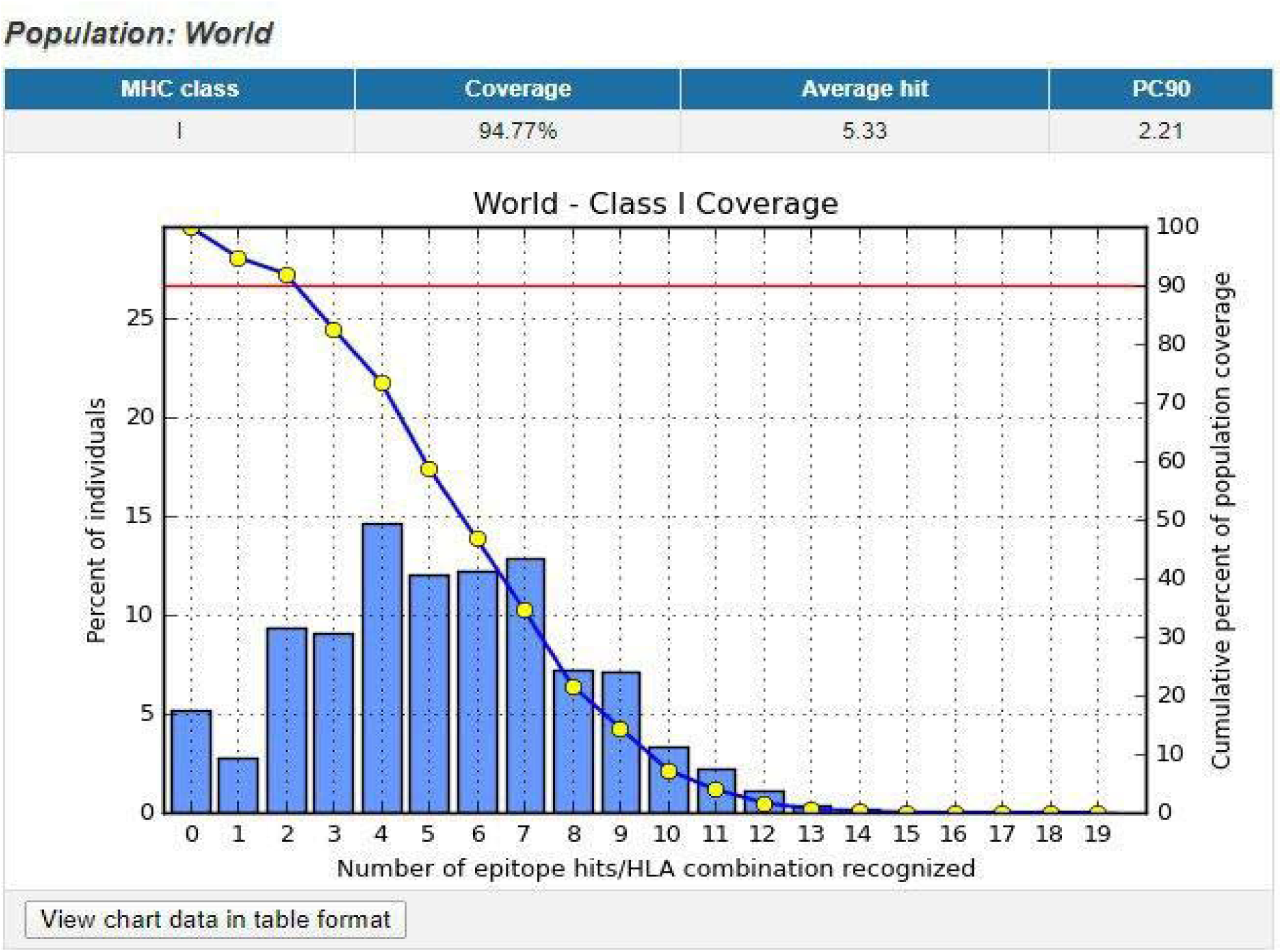
Illustrates the global coverage for the top ten MHC-I peptides (YRYGYTRVA, YGYTRVAEM, YLYRYGYTR, WRFDVKAQM, ALWSAVTPL, LLLQKQLSL, LIADKWPAL, KLFGFCPTR, MYPMYRFTE and FLIADKWPA). Note: In the graph, the line (-o-) represents the cumulative percentage of population coverage of the epitopes; the bars represent the population coverage for each epitope.

#### Population coverage for MHC11

In the case of MHC class II, ten epitopes were assumed to interact with the most frequent MHC class II alleles (KMLLFSGRRLWRFDV, GRGKMLLFSGRRLWR, RGKMLLFSGRRLWRF, GKMLLFSGRRLWRFD, TFTRVYSRDADIVIQ, AVIDDAFARAFALWS, FARAFALWSAVTPLT, MLLFSGRRLWRFDVK, GNQLYLFKDGKYWRF, NQLYLFKDGKYWRFS), inferring a massive global coverage. The highest population coverage percentage of these epitopes worldwide was that of KMLLFSGRRLWRFDV with percentage of 83.74% % (table 6, figure 5).

**Table 6:**
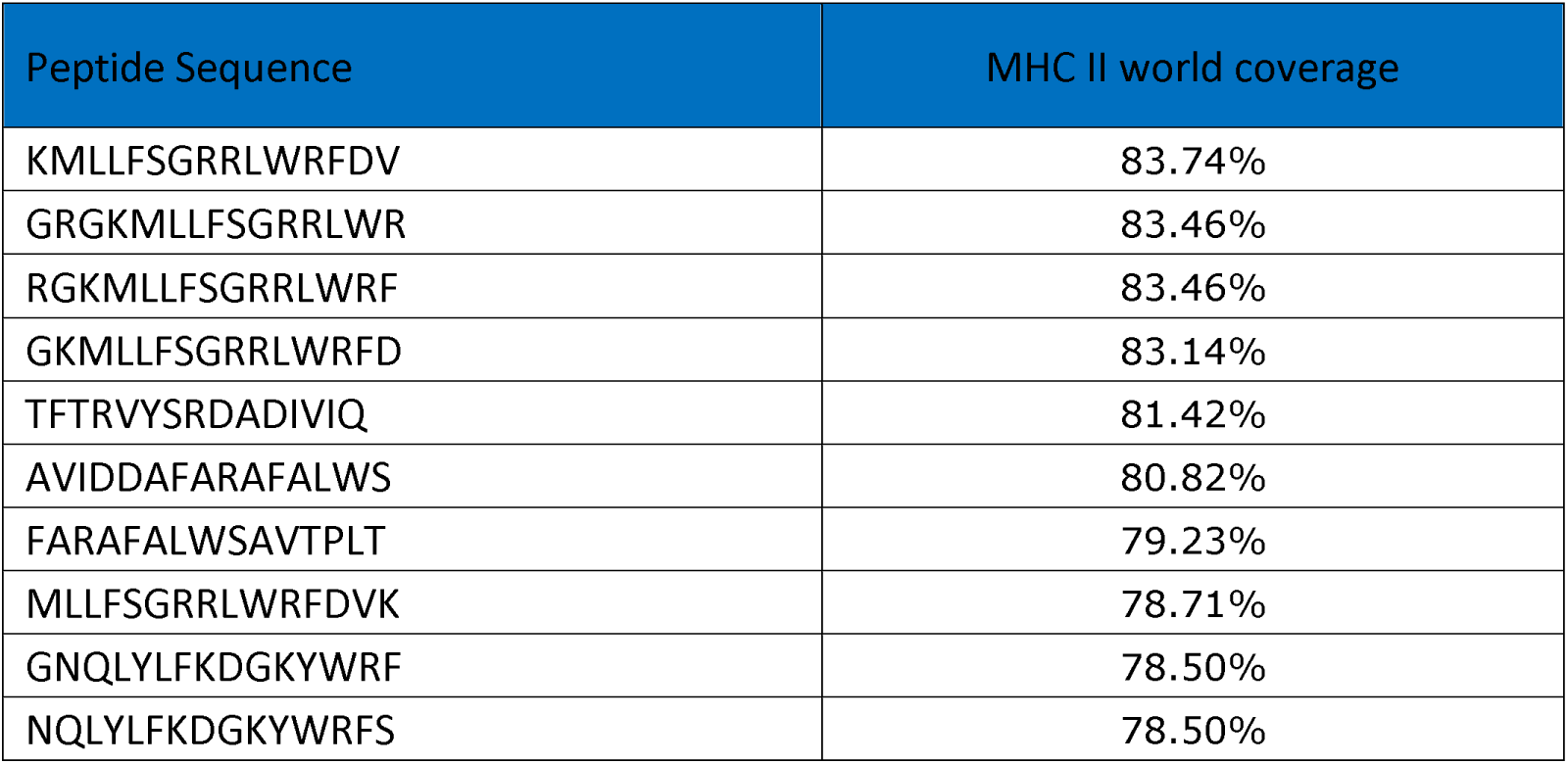
Global Population coverage of promising epitopes in isolated MHC class II.

| Peptide Sequence | MHC II world coverage |
| --- | --- |
| KMLLFSGRRLWRFDV | 83.74% |
| GRGKMLLFSGRRLWR | 83.46% |
| RGKMLLFSGRRLWRF | 83.46% |
| GKMLLFSGRRLWRFD | 83.14% |
| TFTRVYSRDADIVIQ | 81.42% |
| AVIDDAFARAFALWS | 80.82% |
| FARAFALWSAVTPLT | 79.23% |
| MLLFSGRRLWRFDVK | 78.71% |
| GNQLYLFKDGKYWRF | 78.50% |
| NQLYLFKDGKYWRFS | 78.50% |

**Figure 5.**
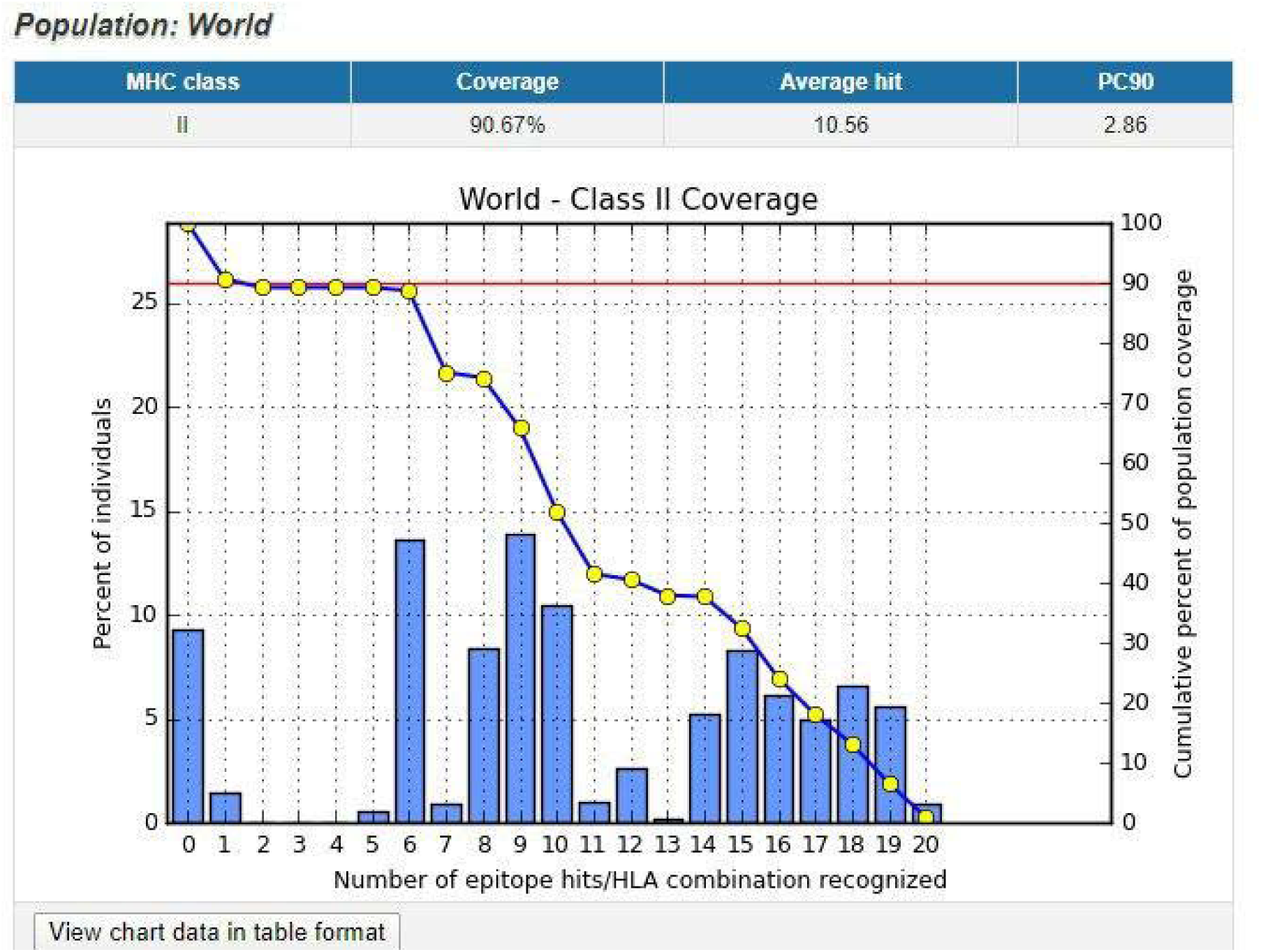
Illustrates the global proportion for the top ten MHC-II epitopes (KMLLFSGRRLWRFDV, GRGKMLLFSGRRLWR, RGKMLLFSGRRLWRF, GKMLLFSGRRLWRFD, TFTRVYSRDADIVIQ, AVIDDAFARAFALWS, FARAFALWSAVTPLT, MLLFSGRRLWRFDVK, GNQLYLFKDGKYWRF, NQLYLFKDGKYWRFS). Notes: In the graph, the line (-o-) represents the cumulative percentage of population coverage of the epitopes; the bars represent the population coverage for each epitope.

#### Antigenicity analysis

Antigenicity analysis of the 10 chosen peptides of MHC1 and MHC2 by vaxiJen software shows four peptides to be probable antigenic peptides in MHC1 (YLYRYGYTR, WRFDVKAQM, LIADKWPAL, and MYPMYRFTE) while only one peptides was found to be antigenic in MHC2 (RRLWRFDVKAQMVDP). (Table 7).

**Table 7:** list of antigenic peptides inMHC1 and MHC2 as predicted by VaxiJen software.

| MHC | Peptide Sequence | Prediction | Score |
| --- | --- | --- | --- |
| MHC1: | YLYRYGYTR | Probable antigen | 0.7645 |
|  | WRFDVKAQM | Probable antigen | 0.7671 |
|  | LIADKWPAL | Probable antigen | 1.2157 |
|  | MYPMYRFTE | Probable antigen | 0.7337 |
| MHC2: | RRLWRFDVKAQMVDP | Probable antigen | (0.5585) |

#### Prediction of interferon-gamma inducing epitopes

IFNepitope server showed three peptides in MHC2 with an ability to induce interferon-gamma production. (GPFLIADKWPALPRK, GRGKMLLFSGRRLWR, and FARAFALWSAVTPLT). Table 8.

**Table 8:** list of MHC2 related peptides that can induce interferon-gamma production.

| Peptide | Prediction | Score |
| --- | --- | --- |
| GPFLIADKWPALPRK | positive | 1 |
| GRGKMLLFSGRRLWR | positive | 0.047075488 |
| FARAFALWSAVTPLT | positive | 0.40888389 |

#### Structure modeling and validation by chimera

MMP9 protein structure was modeled by chimera software and the chosen peptides related to B cell, MHC1 and MHC2 peptides were then visualized.(figure 6,7, and 8). Furthermore, the modeling was verified using the Ramachandran plot in chimera software, which shows that most of the amino acids residues are located the allowed region. (Figure 10).

**Figure 6.**
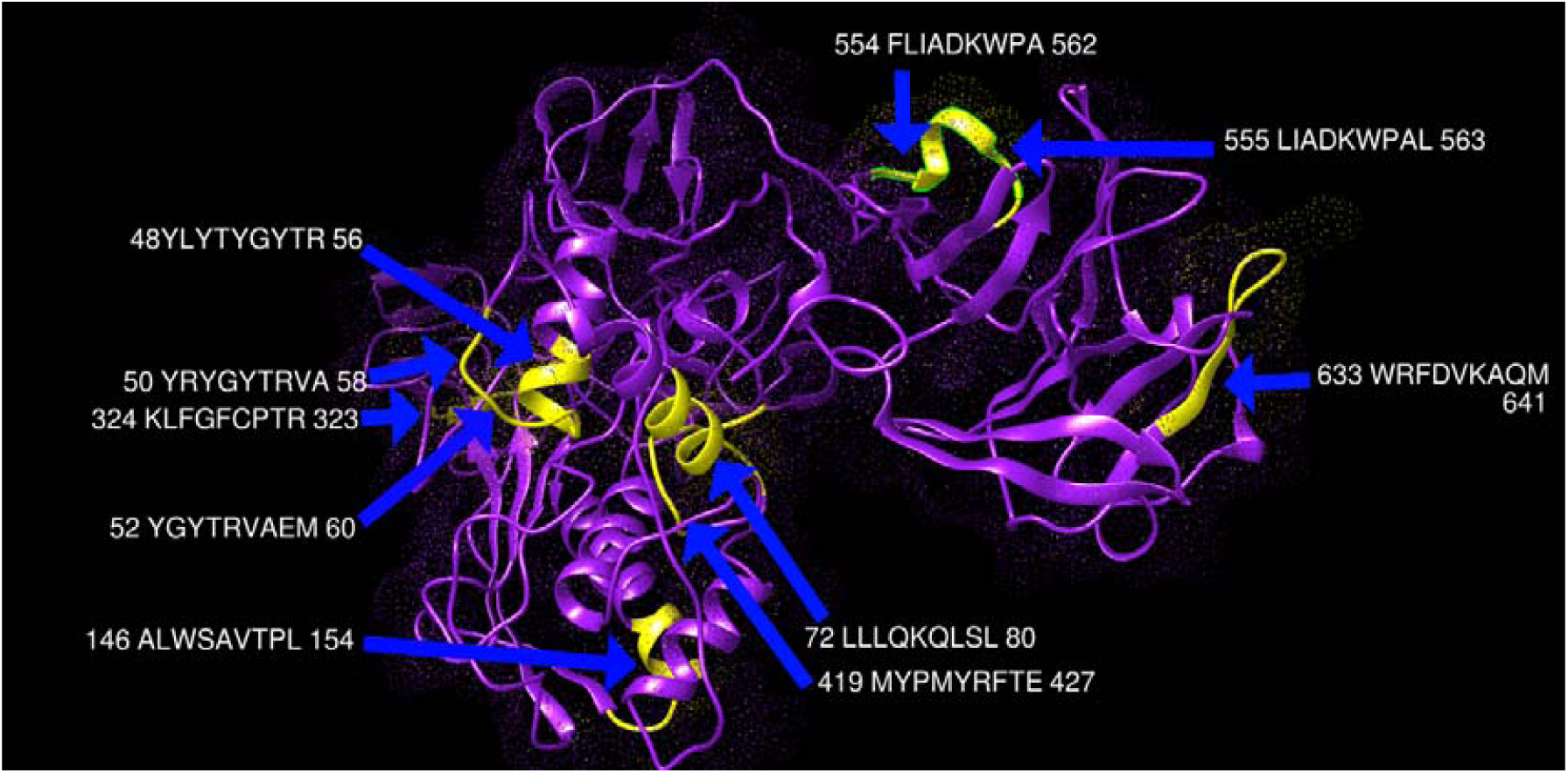
Present 3D structure of metalloproteinase-9 protein visualizing top ten T-cell peptides binding to MHC class I using chimera (version 1.14).yellow color indicate the chosen peptides while the purple color indicate the rest of the molecules.

**Figure 7.**
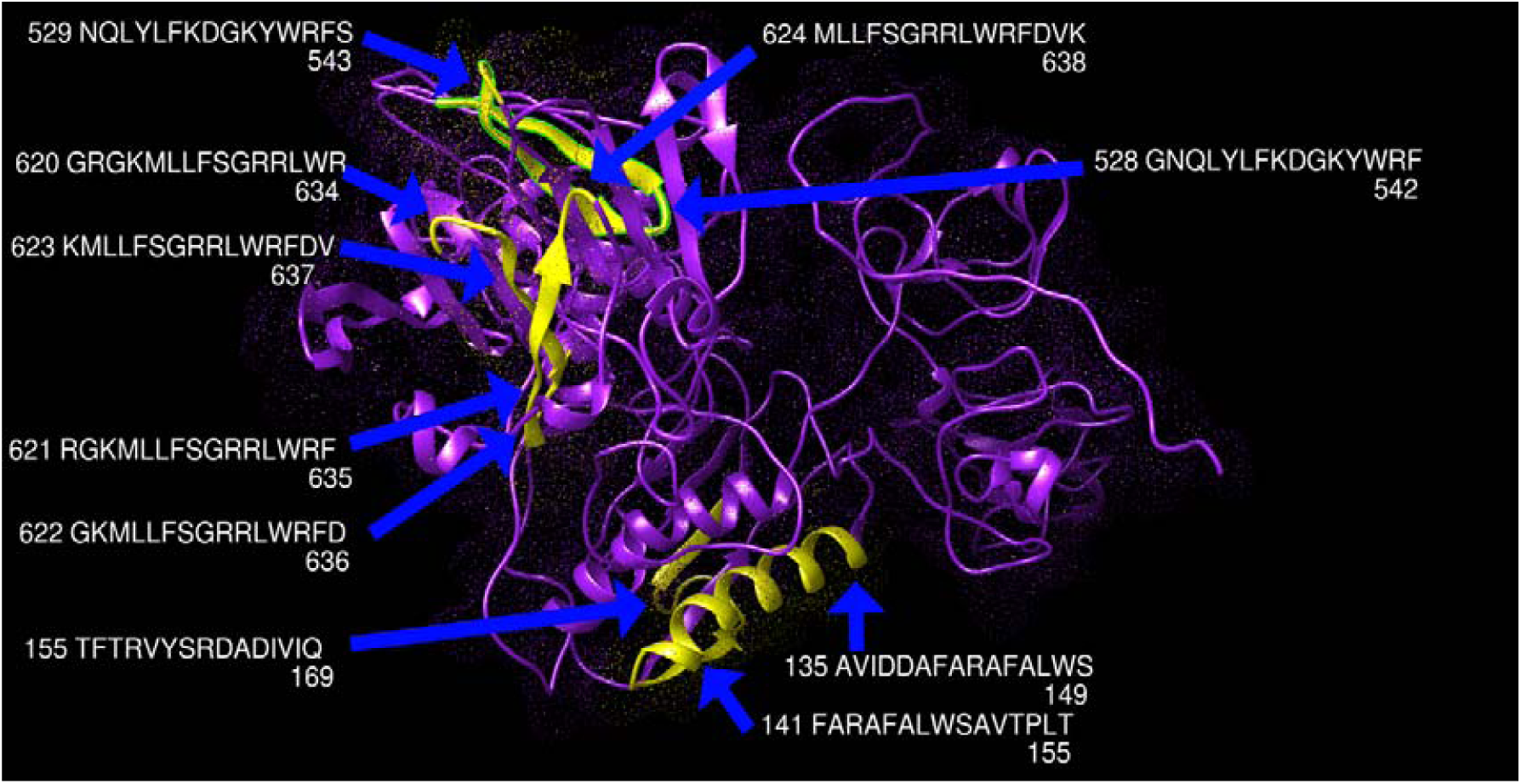
Present 3D structure of metalloproteinase-9 protein visualizing top ten T-cell peptides binding to MHC class II, using chimera (version 1.14).yellow color indicate the chosen peptides while the purple color indicate the rest of the molecules

**Figure 8.**
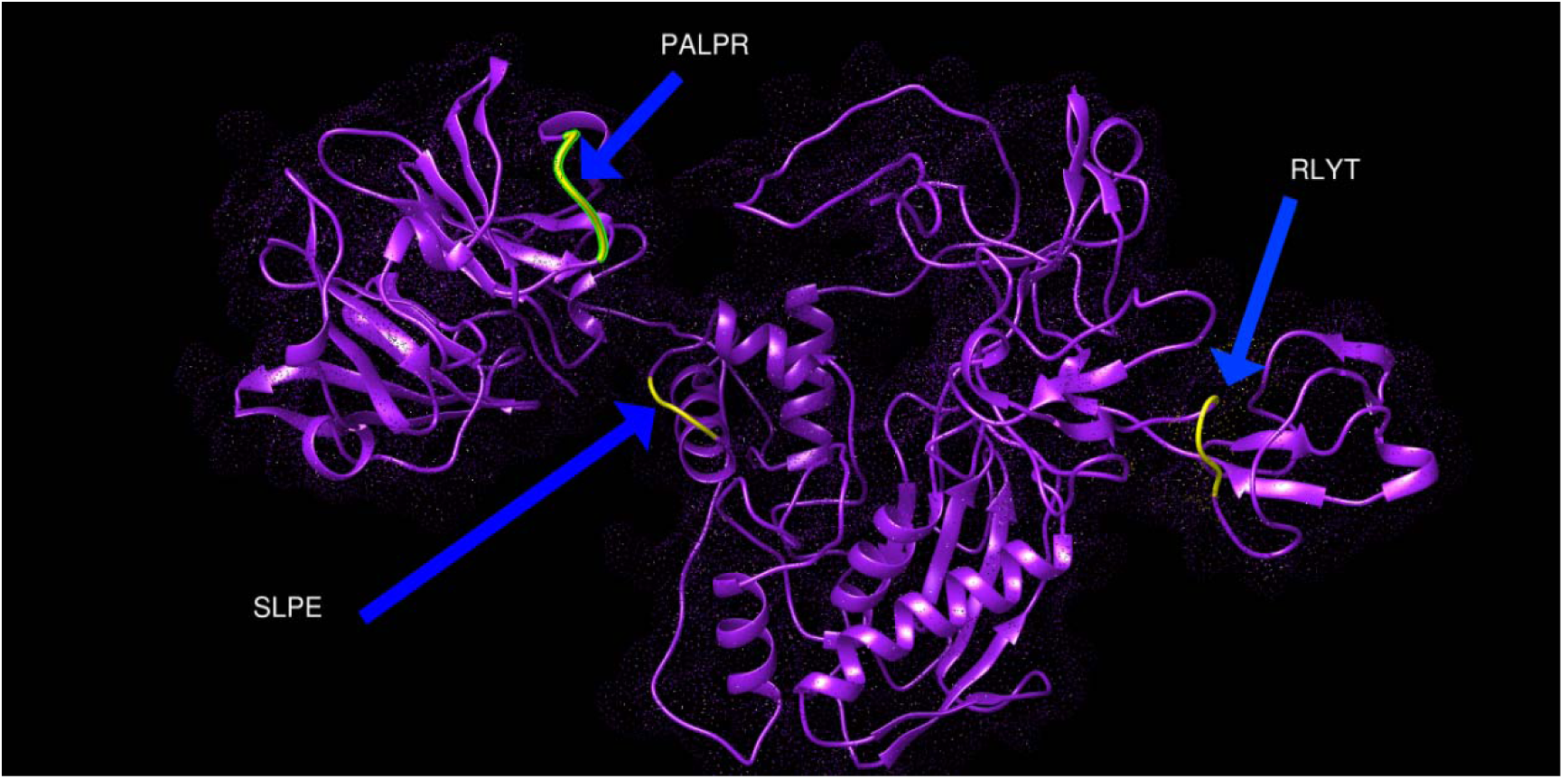
Present 3D structure of metalloproteinase-9 protein visualizing most promisisng B-cell peptides that have high suface accessibility and antigenicity scores, using chimera (version 1.14).yellow color indicate the chosen peptides while the purple color indicate the rest of the molecules

**Figure (10):**
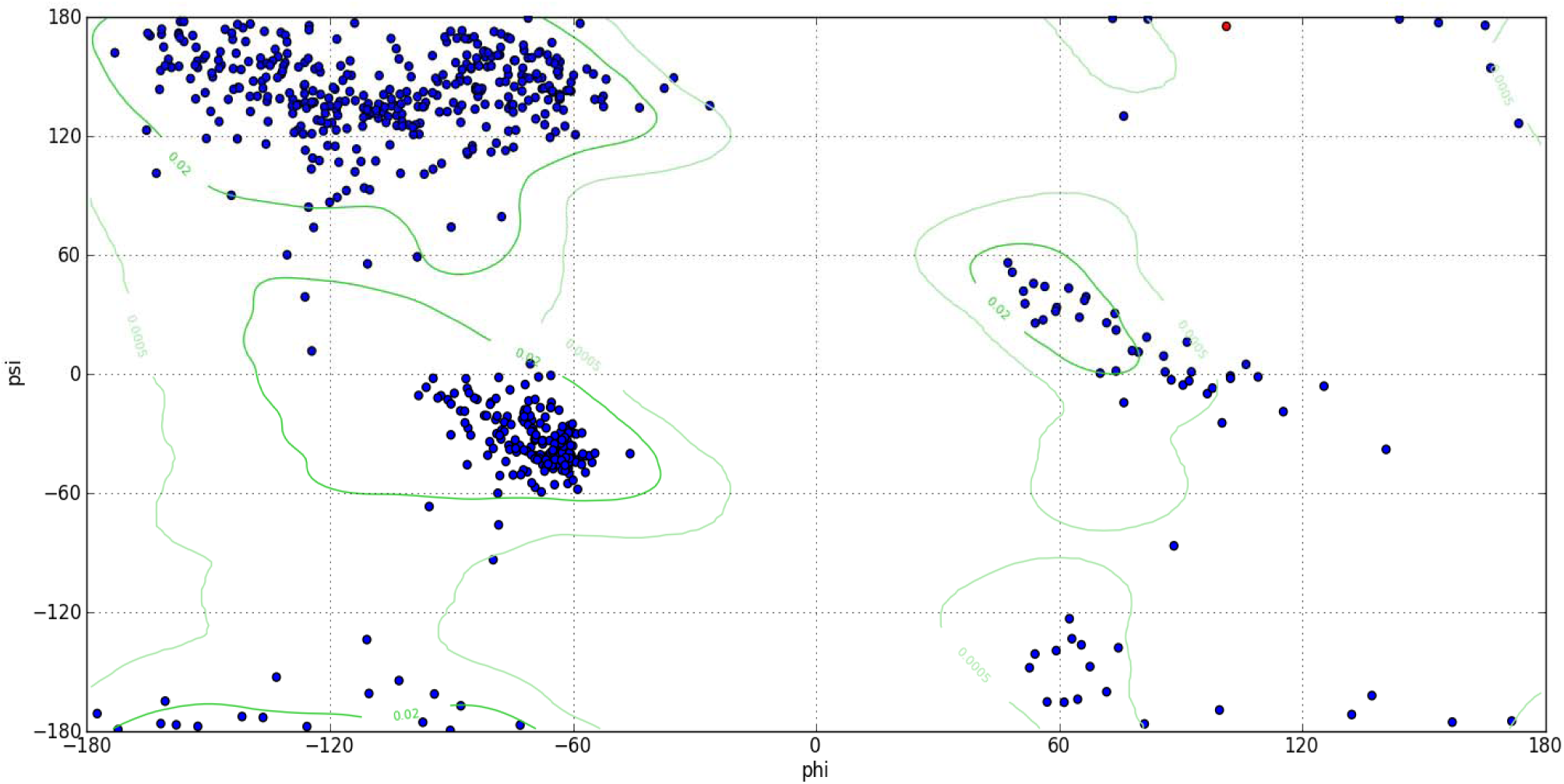
Ramachandran plot analysis of MMP9 protein showing most of the torsion angles located at allowed region (the blue dots represent torsion angles, the green lines indicate the allowed region. (phi) □ and (psi) □ are torsion angles. The torsion angle about the N—C bond is called □ and that about the C—C bond is □. This analysis predicted by chimera version 1.10.2.

**Figure 11:**
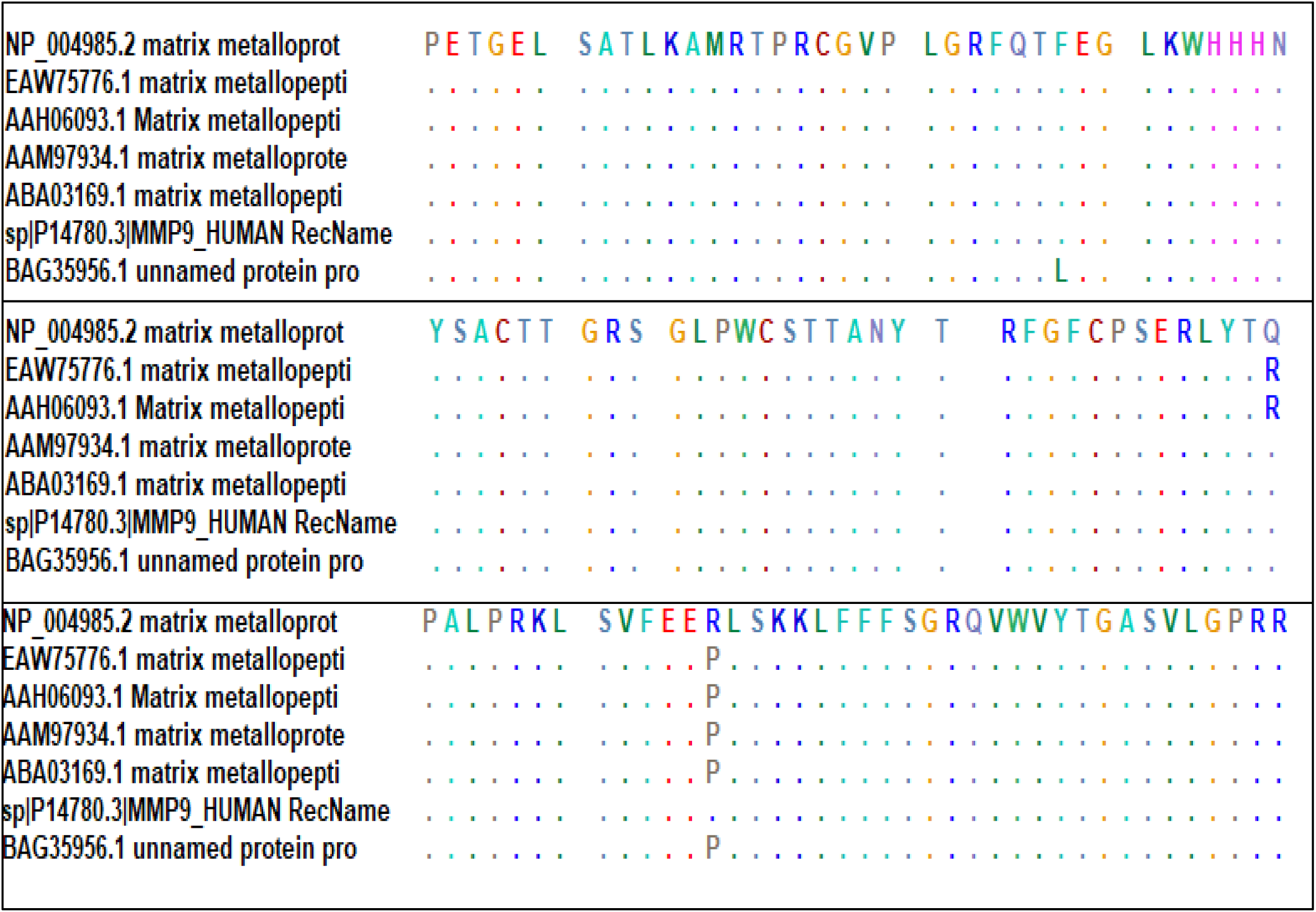
MMP9 sequences conservation test by BioEdit software showing three non-conserved region at positions 110, 279, and 574. The continues colored dots represent conservation.

**Figure (12):**
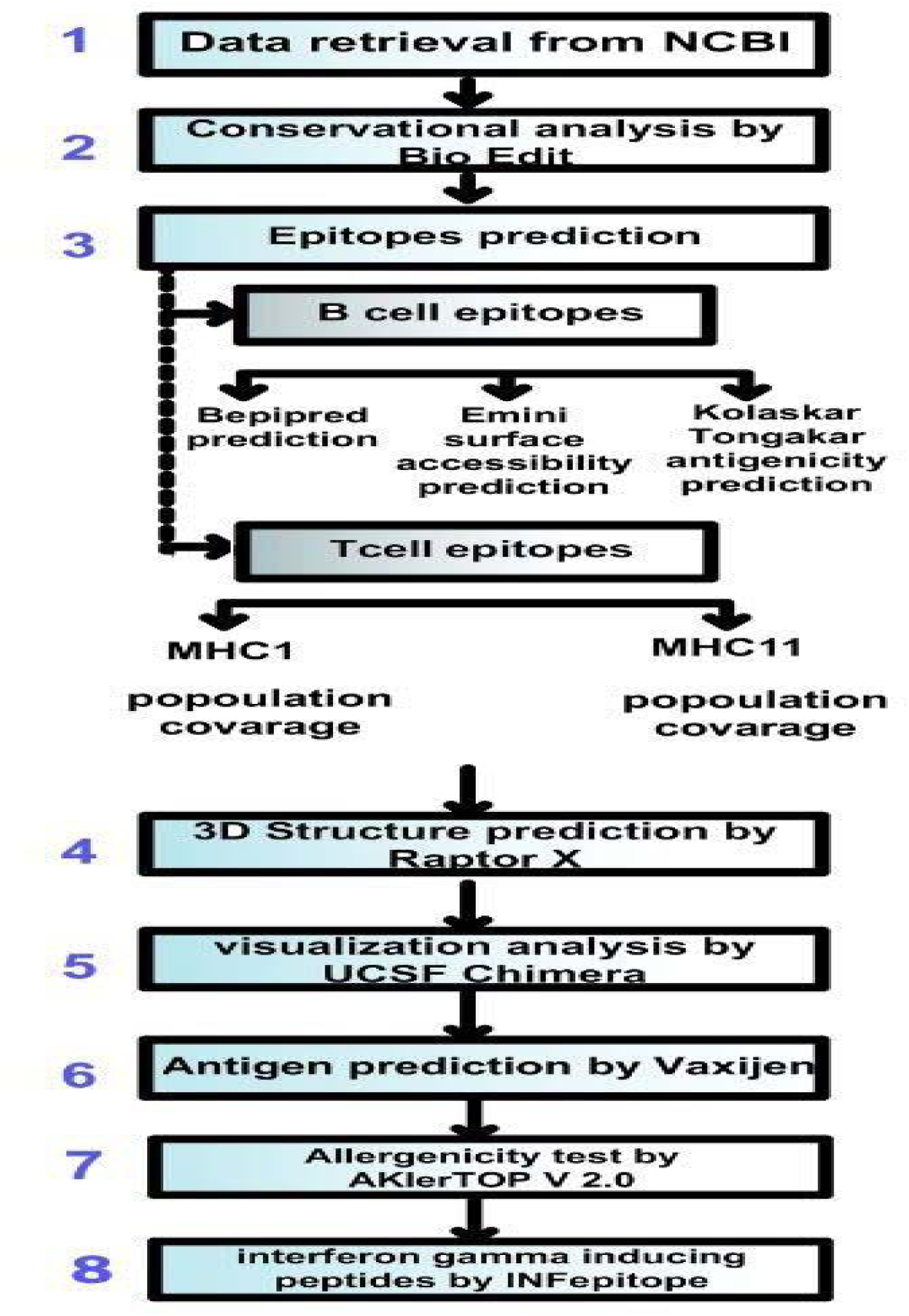
Flow chart of MMP9 vaccine design, showing the different software used for the analysis and design of the targeted peptides.

## 4. Discussion

Multi Epitope peptides will be the next generation of cancer immunotherapy. However, an efficient vaccine progression and production are costly and time consuming, but the development of immunoinformatics tools has made a broader way in developing vaccines through understanding the immune response within a short amount of time.(53) Many research aimed to design universal cancer vaccines using different approaches, yet until now, there is no FDA approved universal cancer vaccine.(54) The current work used immunogenic protein (matrix metalloproteinase-9) that is essential for the survival of all type of cancer as a target for vaccine design, where three Peptides found to be good candidate for interactions with B cells, in addition ten peptides found as a good target for interactions both MHC1 and MHC2. The different soft wares used for the analysis are summarized in Figure (12).

Most cancer vaccines depend on T cell interactions but recent research indicates the significant role B cell plays in the immunology of cancer vaccine, B cell produces antibodies and help in the regulation of other immune reaction.(55) furthermore, several B cell based cancer vaccine studies show Promising results in chronic lymphocytic leukemia and other types of tumors.(56) In the current work, Linear B cell epitopes of matrix metalloproteinase-9 analyzed by Bepipred linear epitope prediction tool. After checking for conservation using BioEdit software (figure 11), the epitopes were checked for surface accessibility and antigenicity by using the Emini surface accessibility prediction method and Kolaskar and Tongaonkar antigenicity method respectively.(Figure 1,2, and 3) The following peptides have the highest score and were approved as the most promising B cell epitopes (SLPE, RLYT, and PALPR). (Table 1 and 2).

T cell receptor can only identify epitopes if it is presented on major histocompatibility complex, Epitopes binds in linear form to MHCs, they linked together into the binding groove of MHC class I and class II through interactions with the amino acids located in the pockets on the floor of MHC.(57, 58)

Cytotoxic T cell(CD8) can only recognize antigen when it is presented on MHC1, and so we analyzed the peptides that has the highest population coverage to and the best binding MHC1 and found 10 candidates(YRYGYTRVA, YGYTRVAEM,YLYRYGYTR,WRFDVKAQM, ALWSAVTPL, LLLQKQLSL, LIADKWPAL, KLFGFCPTR, MYPMYRFTE, FLIADKWPA)(Table 3)(Figure 4). It is interesting to note that among the associated alleles related to these peptides, we found the most frequent reported allele in Caucasians which is HLA-A*0201.(59) This further proves the significant coverage of these peptides.

In the case of T helper cells, it can only recognize antigen when it is presented in MHC2. Therefore, we predicted the peptides of MMP9 that can firmly bind with MHC2, which also hold the highest population coverage and end up with the following 10 peptides sequence (KMLLFSGRRLWRFDV, GRGKMLLFSGRRLWR, RGKMLLFSGRRLWRF, GKMLLFSGRRLWRFD, TFTRVYSRDADIVIQ, AVIDDAFARAFALWS, FARAFALWSAVTPLT, MLLFSGRRLWRFDVK, GNQLYLFKDGKYWRF, and NQLYLFKDGKYWRFS). (Table 4)(Figure 5)

The population coverage of the epitopes and their respective HLA alleles are crucial, as HLA alleles are highly polymorphic in nature.(60) We aim for this vaccine to benefit patient from different parts of the world, therefore, we analyzed the whole word population coverage for our predicted peptide, where we found MHC1 and MHC2 to have population coverage of 94.77% and 90.67% Respectively (Table 5 and 6). Therefor our vaccine is assumed to elicit a strong comprehensive immune response against our targeted protein matrix Metallopeptidase 9 (MMP9) in the majority of vaccinated population.

All peptides were visualized using chimera software (Figures 6, 7, and 8) which shows the predicted structures and positions of these peptides in MMP9 protein(yellow color), then Ramachandran plot analysis shows most of the amino acids residue located in the allowed regions which validate the accuracy of the generated structure.(61)

Allergencity is an important factor in vaccine design as allergic reaction could be fatal to the patients and hence we tested the allergencity of our proposed vaccine using allertop software which confirm the non allergencity of the vaccine.(62) Furthermore the antigenicity is a significant factor to tack into account when choosing a target for vaccine design, (63) therefore we tested the antigenicity of mhc1 and mhc2 peptide using vaxiJen software which shows 4 peptides to be antigenic in mhc1(YLYRYGYTR, WRFDVKAQM, LIADKWPAL, MYPMYRFTE) and one peptide to be antigenic in mhc2 (RRLWRFDVKAQMVD). (Table 7)

Interferon gamma is a cytokine that plays an essential role in the innate and adaptive immunity. It is produced from T cell upon activation by specific peptides which lead to improvement in cancer elimination by inducing ischemia and improving recognition of the tumor cells by CD4 and CD8 cells.(64) we tested the ability of our chosen peptide to induce secretion of interferon gamma through IFNepitope software which indicate that there is 3 peptides in MHC 2 to positively affect the interferon gamma production (GPFLIADKWPALPRK, GRGKMLLFSGRRLWR, FARAFALWSAVTPLT)(Table 8).

In a study done by Roomi et al, where they used MMP9 as anticancer vaccine in mice, they found 80% reduction in tumor volume after using this vaccine.(65) Which further proves the predicted effectiveness of our proposed vaccine. We took further step in our work to analyze this target using human database, which confirm our hypothesis about the role of MMP9 as a good candidate for vaccine.

Previous research by Dosset et al. designed a universal peptide based cancer vaccine using telomerase reverse transcriptase (TERT) which showed improve response against CD4 and CD8, but the role of B cell was not investigated in this research.(66)

This study is limited due to the few numbers of reference sequence we were able to acquire from NCBI, along with the 6 alleles(HL-DRB3*03:01, HL-DRB3*02:01, HL-DRB4*01:03, HL-DRB3*01:01, HL-DRB4*01:01, HL-DRB5*01:01) that was not applicable for prediction in IEDB.in addition to the limited therapeutic activity of peptide vaccine in general.(67)

In the end, 23 peptides were chosen as a target for universal cancer vaccine design, yet Wet lab studies are required to confirm these results, in order to move another step towered the goal of eradicating cancer from the face of the earth.

## 5. Conclusion

The current work designs multi epitopes peptide based universal cancer vaccine using MMP9 protein as a target where three Peptides found to be good candidate for interactions with B cells, and ten peptides found as a good target for interactions with each MHC1 and MHC2.

## 7. Acknowledgement

The authors acknowledge the Deanship of Scientific Research at University of Bahri for the supportive cooperation.

